# Exploring Intuitive Approaches to Protein Conformation Clustering Using Regions of High Structural Variance

**DOI:** 10.1101/2021.09.05.459014

**Authors:** Yi Yao Tan

## Abstract

This paper presents a method to find structurally high variance segments of the different conformations of a single protein and uses clusters them using different distance metrics and interpretation of coordinate and angle data presented by three different methods: root mean squared derivation RMSD, t-distributed Stochastic Neighbor Embedding (t-SNE) based map, and dihedral-based clustering. The methods were applied on the human cylin-dependent kinase 2 (CDK2) protein, code P24941 uniprot using a series of python scripts and clustering packages. We test our methods on the data of the CDK2 protein as it is a highly researched protein, with practical applications of clustering in cancer research, crucial in the regulation of the cell-cycle, and has a sizeable amount of experimental data collected on the confirmation structures.

While using the distance based root mean squared deviation RMSD provides data of structure to structure dissimilarity between different conformations, a simple RMSD matrix lacks to ability to describe the subsequence-wise in shape and absolute position which could be the main identifying elements for a protein’s conformation and state. To make up for this loss we explore an intuitive and more flexible method, able to accept multiple high structural variance segments, which takes coordinate based data, through a series of maps and with the help of t-SNE, and maps each segment as a feature in the clustering matrix. This method, however, would require additional testing on other proteins and modifications to verify its consistency and test its robustness.

In the end we explore the pros and cons of the three methods applied on the high structural variance regions. Despite the randomness factor by the t-SNE used in mapping the coordinates to lower dimensions, the coordinate-based approach consistently performed better than the RSMD and dihedral based methods in clustering the three groups of the CDK2 protein kinase. We also found that analyzing only the substructures identified by the high variance detection algorithm consistently provided more distinct clusters with higher multi-class F1 scores.

## Introduction

A lot of different methods and algorithms have been devised to optimize protein clustering including the use of domains of topology and geometry to identify similar protein structures. The uses of this field includes that of notably structure based drug design and other structure based biological research, a wide range of automated clustering algorithms have been developed to facilitate the process in the industry. To address this hierarchical clustering is used to group similar proteins. It relies on the input matrix a derived by a specific and adequate distance metric to identify the conformation.

In this specific experiment we analyze CDK2, which is an enzyme that regulates the G1 to S phase transition or simpler said growth the DNA replication phase. When genetic material is damaged, DNA replication needs to be controlled to avoid flawed replication which is a characteristic of cancer. To address this control this growth, synthetic inhibitors can be developed or natural inhibitors can be used, for example the tumor suppressor p53 (Bartek and Lukas 2001), to prevent the advancement into the S phase. The development of automated clustering techniques can speed up this process of the development of synthetic inhibitors which are then used for medical treatments.

Beyond the application of the specific protein, structural difference in proteins can lead to functional differences between the proteins, by clustering similar proteins scientists might be able to guess the functions or states of newly discovered proteins by structural analysis. Due to the fact that similar protein conformations might share similar structures but not share the same state or function, an analysis can be performed on the structures of the subsequences to identify the key characteristics of each group of proteins or conformations.

In order to amplify these differences in substructures and form clear clusters, we develop a method to identify segments of the sequence with high structural variance between conformations apply the three methods on these specific substructures to cluster the conformations. By applying maps like the RMSD and the later on proposed t-SNE based map on the coordinate information, in particular that of the identified high structural variances segments, we create more homogeneous and distinct clusters for each type of conformation of the protein.

## Method

### 2.1 Data Annotations

The data list of samples for the human CDK2 protein are first retrieved from Uniprot, code P24941 (*UniProt* n.d.) and with a basic shell script a total of 420 pdb files entries of different pdb files are curled from rscb.org (**rcsb**). Then using a python script the pdb files are read and annotated storing the name of the entry, each chain, the resolution, presence of a cyclin structure is used to determine the close and openness of the the individual chains of each of the file. Also, the phosphorylation of threonine (TPO) at position 160 of the chain to determine the activity. In particular, the presence of cyclin means that the chain is open and otherwise closed; and the phosphorylation of threonine means the conformation of active and without— inactive. The scripts then store the annotated data chain by chain then stored into a dictionary with the keys as the name of the chains and values as objects with each of the properties stored within. A total of 531 annotated chains are registered.

### 2.2 Structural Alignment

To begin the structural alignment, a python script is used to pick the chain with the highest resolution and the presence of the natural ligand: ATP in the chain. This choice is then used to align the rest of the chains onto since the highest resolution will give the most accuracy in alignment and since other synthetic ligands, if any, in the other conformations are made to imitate the natural ligand make it a good candidate for the rest of the samples to align to. In the case of the experiment 4EOJ chain A was found as the best choice to align the rest of the conformations to, with a resolution of 1.65 angstroms and the natural ligand of ATP.

Structural alignment uses the software pymol built in function *align* and iterates through all of the loaded chains and aligns the structures with structural outlier rejection cycles set to the default: 5, rejection cut off default of 2.0 and target = 4*EOJ*_*A*. This alignment also transforms the structure coordinates to be placed on top of 4*EOJ*_*A*. All of this is done smoothly since the samples have consistent numbering.

### 2.3 Finding structurally high variance regions

The main idea of the algorithm is to find the high variance segments, in terms of the structure. Using this region, for the Diheral and RMSD methods, a distance matrix is created from analyzing the subsequence with the highest variance. For the t-SNE based method that we can use *n* distinct high variance regions chosen to identify the conformations which will then be mapped to a single value for each of the conformations used as a feature, thus a *number of samples × n* sized matrix.

To define a useful definition of variance of the structure, just by using the variance of coordinates is insufficient to give properties like shape and structure. To trace the structure, vectors from the coordinate of the alpha carbon from one residue are drawn to the next. Theoretically, unaltered bond angles and lengths between the residues of corresponding indices of different are parallel and of the same length, thus vector subtraction will give a result near 0; therefore, by using the variance in the defined vectors a variance in shape. Of course the change in a previous bond angle could affect the position of the next; however, in agglomeration the goal of finding a high variance segment is achieved. More specifically the variance of the *residue_k_* to *residue_k_*_+1_ vector is defined by the average of the variance of the x, y, and z coordinates.

For example the close up of the activation loop of: 1HCK_A in Pale Green —Closed inactive, 1QMZ_A in Sand —Open active, 4EOJ_A in Deep Salmon —Open active:

**Figure 2.1:**
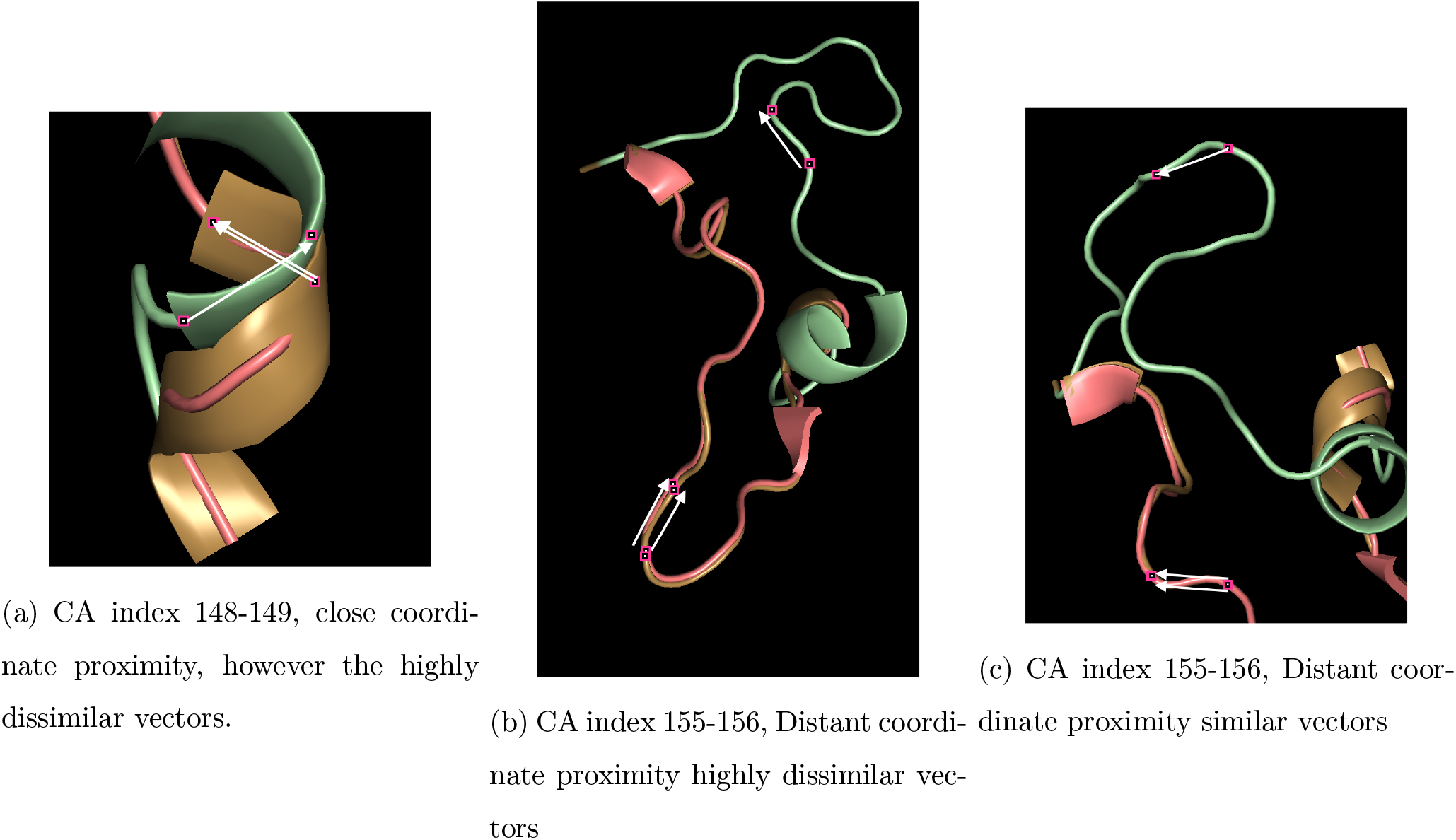
As we can see, the two open active strutures have consistently parallel vectors from residue to the next residue while the closed inactive, 1HCK chain A, often is pointing in a different direction. By a calculation of the average x, y, z component of variance, we can identify regions which would be different structurally.

**Figure 2.2:**
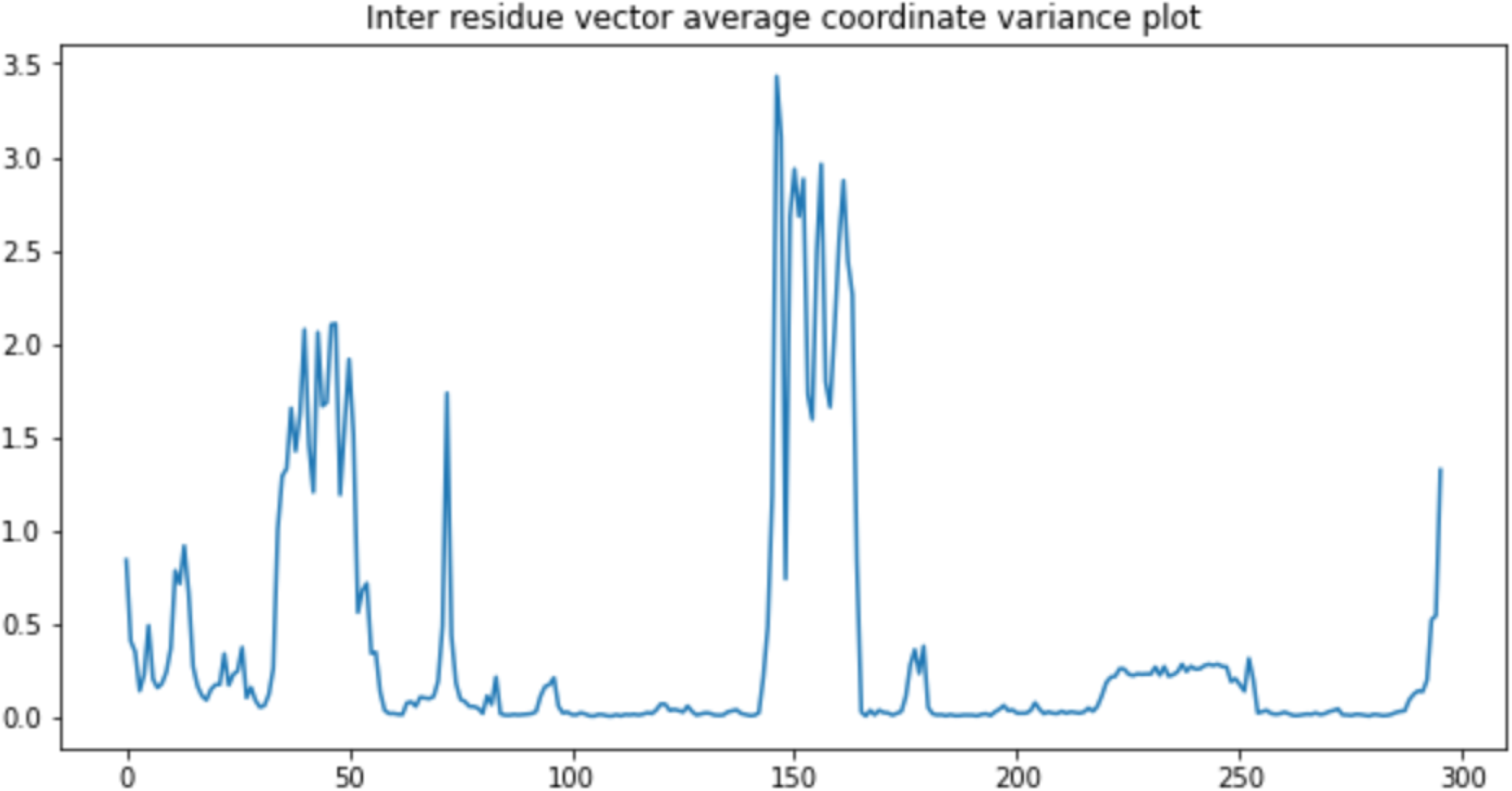
plot of average x, y, z component variance of residue to next residue. Eg: index 0 is vector from CA residue 0 to CA residue 1 (0 indexed)

To gain proper information of the choice of high variance regions, a function is used to obtain all of the indices with above 90-th percentile variances, then the indices are grouped into subsequences which are ranked by their average variance value from highest to lowest in order are: vectors indices 144 to 164 the activation loop with an average of 2.161 *ångströms*^2^, 34 to 54 around the PSTAIRE helix average 1.467 *ångströms*^2^, 71 to 73 with *ångströms*^2^, index 0 0.841 *ångströms*^2^, 11 to 14 with 0.769 *ångströms*^2^ near the glycine rich loop, 293 to 294 with 0.532 *ångströms*^2^.

Now using these high variance segments, we can better identify the coordinate based differences that were previously suppressed by similar regions between all conformations.

### 2.4 RMSD and Coordinate Data

In the same order that the chains are loaded and aligned, a 531 by 531 RMSD matrix is calculated in the same order both in columns and rows the root mean squared deviation of the alpha carbon atoms of residues present in each pair of structures given by:

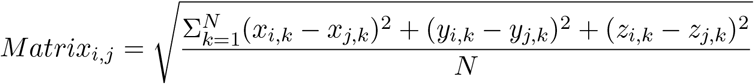

Where *x_i,k_* is the x coordinate in the i-th chain of the k-th residue (out of the total residues present in both chains) and N is the total residues present in both chains. Very clearly, the diagonals of the matrix are all 0s since there is no dissimilarity between a conformation and itself. Lastly, coordinate matrices of each conformation is also obtained by pymol’s *getcoords* command, using python’s module multiprocessing for optimization (Schrödinger, LLC 2015).

### 2.5 Dihedral Based Clustering Technique

Dihedral based clustering is a method implemented by Protein Structural Statistics. In this paper we compare our results to the dihedral based clustering from a paper by Thomas Gaillard. The method uses a distance matrix with the a function of the distance metrics defined by the circular distance, where T is the period. The metric measures dihedral angles within residues which provide an insight to structure. The distance of the k-th dihedral angle of the i-th and j-th conformation is given by:

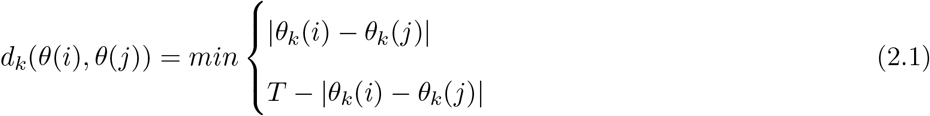

and the distance matrix is defined by:

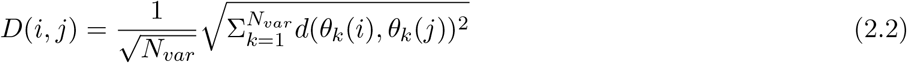

(Gaillard et al. 2013)

### 2.6 t-SNE based map clustering technique

#### 2.6.1 Mapping coordinate to usable feature values

This method doesn’t use a distance matrix and keeps the relative position of each segment (instead of distances) via a map; this allows for clearer distinction between clusters.

Let m be the number of samples, n be the number of feature segments and, *l_i_* be the length of the i-th feature segment. First to make the matrix with missing residue coordinates all usable, in the case where conformations with missing residues need to be classified, sklearn *IterativeImputer* is applied upon a matrix of the arrangement of same index residue coordinates from all of the chains, size of *m ×* 3, in this case since if a residue is none all of its coordinates are not available, Multivariate Imputation by Chained Equations is not able to help fill in the row of the matrix, thus *IterativeImputer* actually imputes using the average of x, y and z of the coordinates of the specific index of the chain over all of the samples. This process is repeated for every index of the subsequence size of *l_i_*. After the imputations are calculated, the coordinates are then arranged back by samples.

A map is needed to send a matrix *l_i_ ×* 3 of coordinates of each high variance segment to a single value so that the values can be used as features to represent the conformation in clustering. Since in this region the coordinates vary a lot of and the shape too, it makes sense to find the average of the coordinates and perhaps similar segment structures would cluster together having similar arrangements of coordinates.

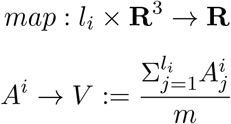

Where 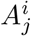 is the coordinate of the j-th index residue of the i-th variable segment. This results in a *m ×* 3 matrix for each high variance segment where m is the number of samples and each sample has an (*x, y, z*) average coordinate value. Now using the fact that similar structures are already clustered in three dimensions, to maintain cluster distances on a projection to a one dimensional vector of values for all of the conformations of the specific segment, t-Distributed Stochastic Neighbor Embedding or t-SNE from the Sci-kit-learn tool kit is used to project the distributions to lower dimensions while preserving inter-clusteral distances giving a final *m ×* 1 vector for our specific segment. This process can then be iterated *n* for a choice of *n* high variance segments depending on the user.

t-SNE constructs probability distributions centered around each point given standard deviation derived by user input perplexity. The similarity between the reference point and another is a conditional probability value derived from the probability density beneath a Gaussian distribution centered at point of reference. Once the probabilities are defined they’re projected into a lower dimension under with a student-T distribution while minimizing the Kullback-Leiber divergences between the probabilities on each dimension which penalizes on further distances between points therefore prioritizing local structures making t-SNE a natural choice for reducing 3 dimensional clustered coordinates into single values. For the sake of creating more distinct clusters and facilitate hierarchical clustering, the value of perplexity used is 50 the highest value that the paper recommended, however, different perplexity values can be chosen depending on user (Maaten and Hinton 2008). Finally random initialization is set to PCA for consistency in results. The t-SNE function first projects data using PCA with a randomized svd_solver and then runs gradient descent of the KL divergence on the PCA projection to create the final embedding matrix (Pedregosa et al. 2011).

The value vector from each segment can be put together in finally creating an *m × n* matrix where m is the number of samples and n is the number of featured segments. Now the *m × n* matrix is ready for clustering. When plotted on the final matrix of features conformations with similar substructures on certain segments will cluster together on each column, and it can be used to describe well the physical similarities of each structure relative to another. Due to the random initialization of t-SNE, to obtain the final results, we run multiple trials and find the average. To find the final prediction of a conformation, we can use the most frequent prediction of the conformation across the multiple trials.

### 2.7 Clustering

The distance matrices are then put through a scipy’s hierarchy clustering, a python package (Virtanen et al. 2020), using ward’s algorithm which minimizes the variance in the cluster after merging. For the purposes of the experiment the dendrogram is automatically cut into 3 using cophenetic distances by sklearn.cluster’s *AgglomerativeClustering*. The result is then plotted using t-SNE projections and analyzed with multi-label statistics. Specifically, using the same definitions:

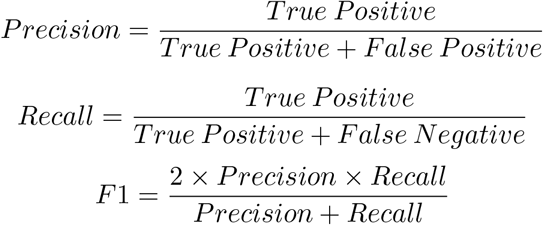

For example we have a 3×3 cross classification table M numbered in the same order of rows and columns, the diagonal will be the True Positives. We define the labels of columns as “real *group-name*” and the rows as “predicted *group-name*.” Thus for precision of *M_ii_* we’ll have:

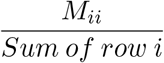

and for recall we’ll have:

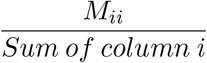

And finally the F1 score will be calculated in the same way. Finally the average across all the 3 groups’ precision, recall, and F1 scores are displayed.

### 2.8 Statistics

After the tree cut, a function tests the permutation of the three clusters to fit the three annotated group, the permutation with the most amount of correct predictions is used to access the F1, precision, and recall score using multi-label evaluation metrics.

For the t-SNE based map clustering technique, 20 trials are used, and then the average is found of the 20 trials of each of the F1, Precision, and Recall for each of the groups metrics and then divided by the number of iterations and similarly done for the the cross-classification tables.

In the end the RMSD clustering, t-SNE Based Map clustering, and dihedral based clustering based on the paper (Gaillard et al. 2013) are compared.

## Results

The results^1^ of only conformations with no missing residues for the full sequence without isolating high variance regions of each of the methods are not shown since only 55 out of 531 have no missing residues. This sample size under represents the total data set. RMSD had a resulting F1 score of 0.396 on the set of 55 conformations while t-SNE based map clustering had an average F1 score 0.445 on 20 runs.

In this section we first explore the results of running the RMSD on all the non-missing alpha carbons between all possible pairwise combinations of two conformations of the data set. This experiment is done on the complete data set no matter the amount of missing residues and RMSD is applied on the whole sequence, provided that there is the residue at the index is not missing in either one of the conformations. Then we explore the RMSD of CAs of all conformations without missing residues in the activation loop which we previously identified as a substructure with the highest average structural variance.

To compare the results, we run the t-SNE based map clustering on the complete data set and of the whole sequence. Additionally the t-SNE based map clustering is also run on the activation loop of all the conformations without missing residues.

Finally we run dihedral based clustering on the activation loop of all the conformations without missing residues; thus we’ll have three methods with identical inputs of the activation loop which we’ll able to compare their performances head to head.

### 3.1 Clustering RMSD results

#### 3.1.1 Complete RMSD CA

The first experiment was done on the complete RMSD^2^ matrix of all the conformations and of the whole sequence.

**Figure 3.1:**
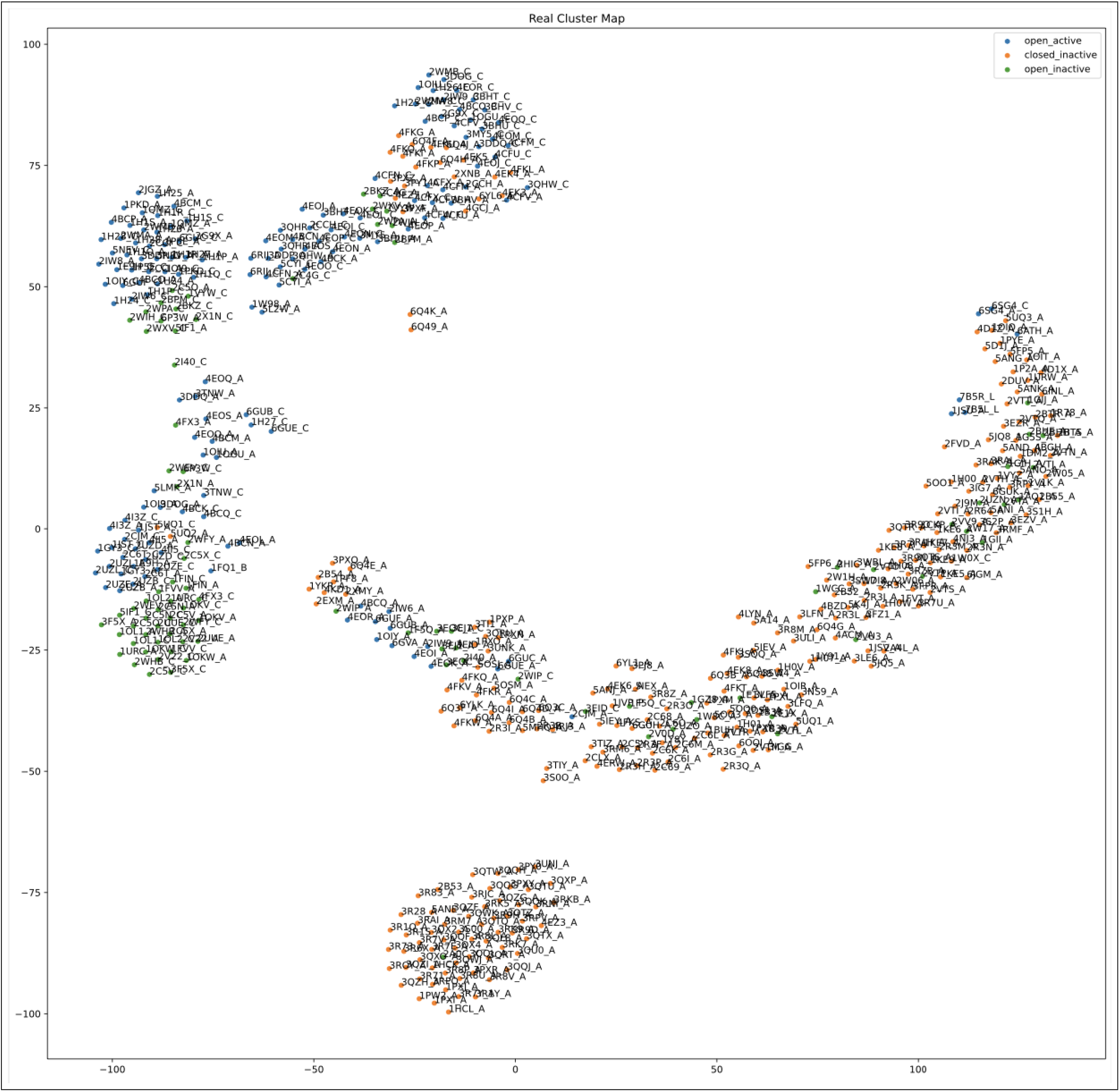
t-SNE projection map of the real clusters.

**Figure 3.2:**
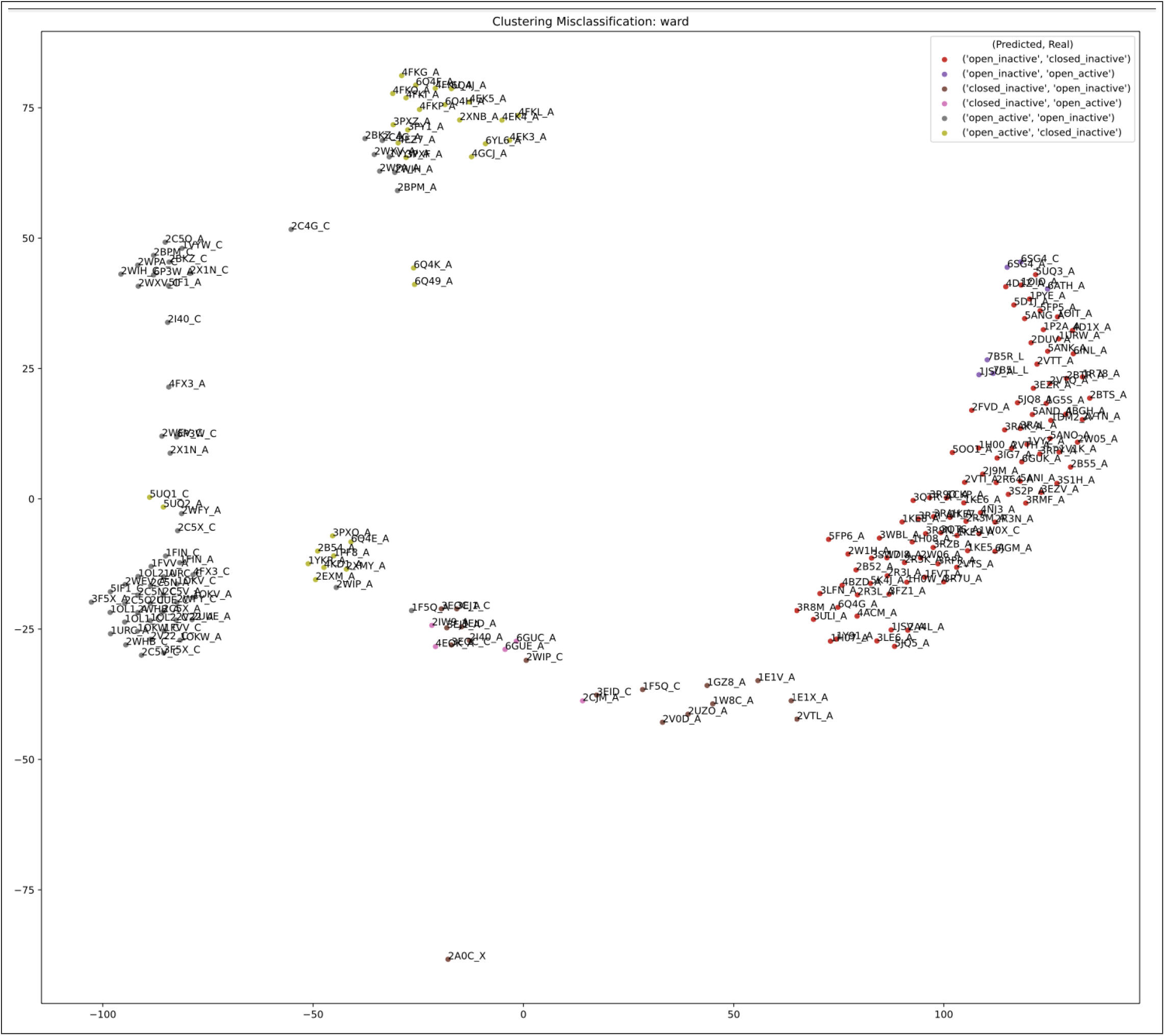
Misclassification Clusters map

**Figure 3.3:**
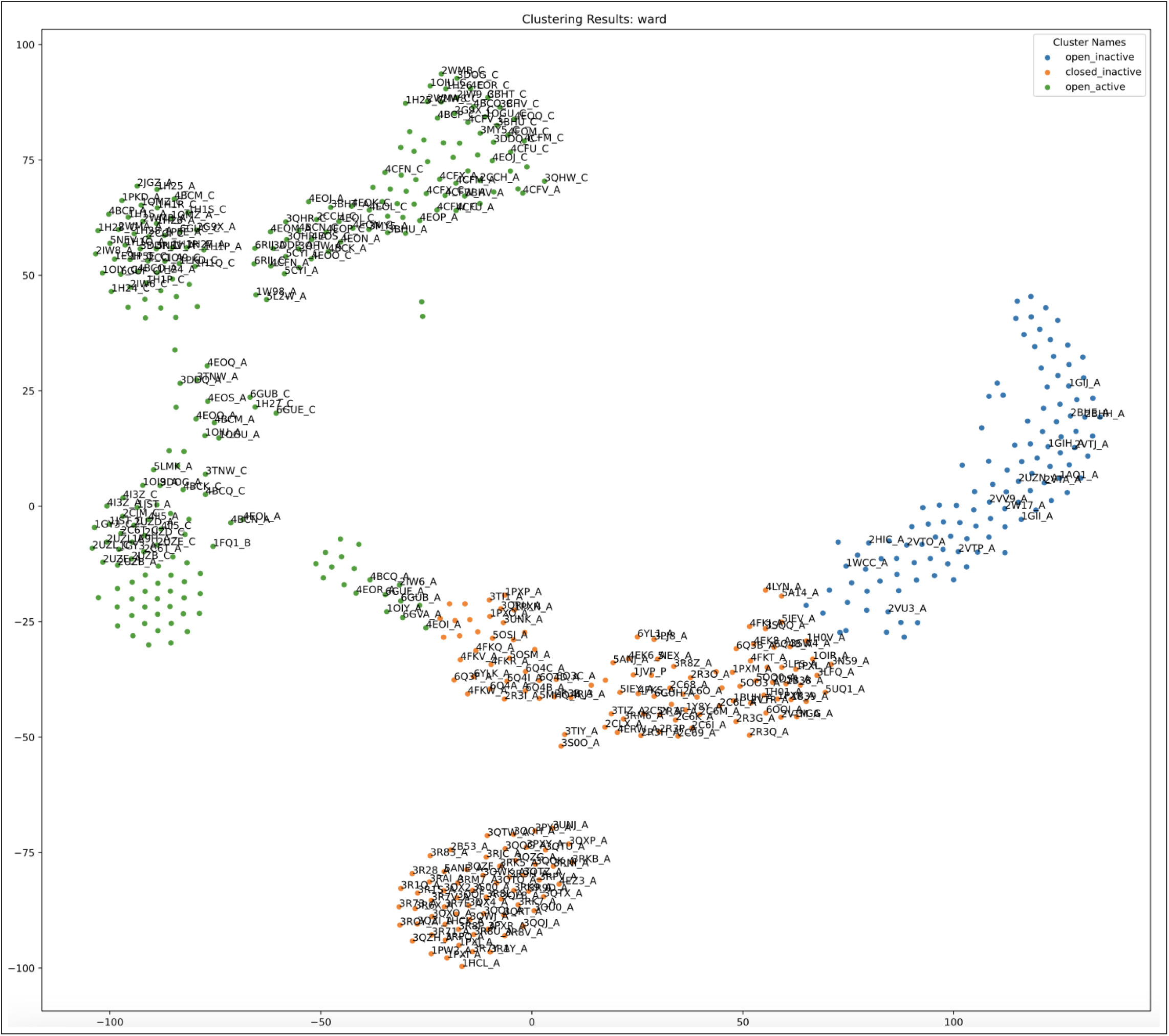
Clustering Map: the correctly classified are labeled and incorrect, unlabeled.

**Figure 3.4:**
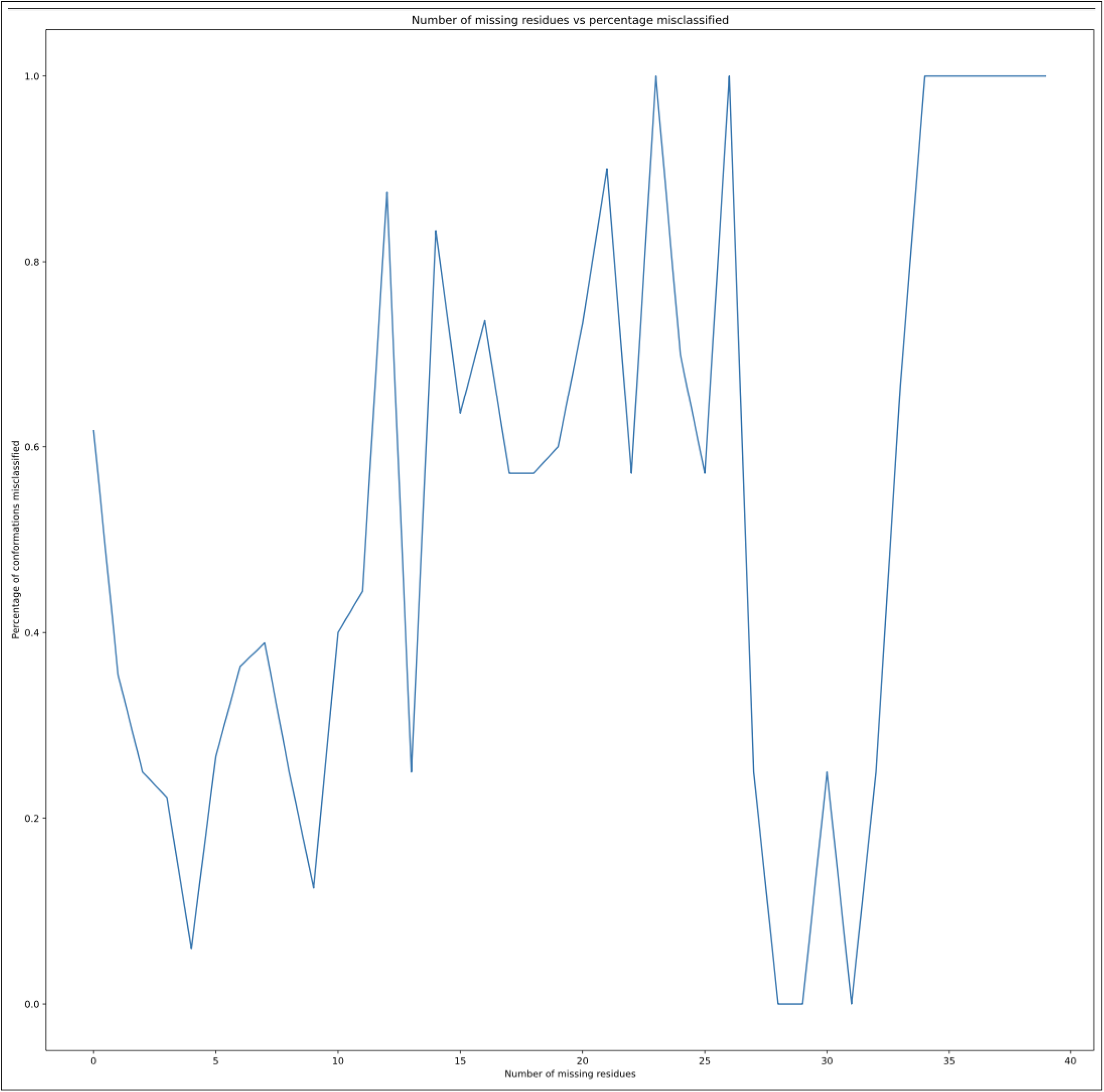
Missing residues vs Percentage misclassified

##### Statistics

**Table 3.1:**
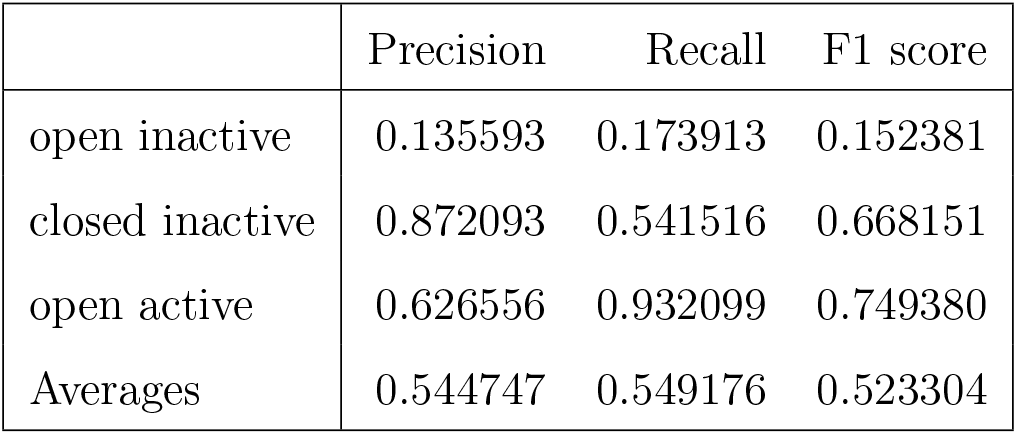
Complete RMSD Activation evaluation metrics

**Table 3.2:** RMSD no threshold of missing residues, full sequence, Cross Classification Statistics

|  | Real open inactive | Real closed inactive | Real open active |
| --- | --- | --- | --- |
| Predicted open inactive | 16 | 96 | 6 |
| Predicted closed inactive | 17 | 150 | 5 |
| Predicted open active | 59 | 31 | 151 |

##### Analysis

From just the t-SNE projection of the real groupings based on the annotations, we can see that first of all, it is hard to identify three distinct clusters. Secondly some of the clusters, especially the one in the top left of the real cluster map is heterogeneous.

Open inactive having the some of the qualities of the two other groups is mostly misclassified as the real open active, while closed inactive is classified often as open inactive. Indeed the RMSD does a much better of separating completely different conformations between open active and closed inactive. Perhaps if the clustering requirements only asked for two different groups RMSD would perform much better, however having one group with mixed qualities, RMSD failed to distinguish the differences. In general the full RMSD doesn’t give very distinct patterns and clusters.

Final it can be seen that with RMSD matrix of the CA data, there is not much of a correlation between the missing data and the misclassification, thus making it hard to identify the missing residues as a factor of mis-classification in incomplete data. A surprising 60% of the conformations with 0 missing residues were misclassified. Either the conformations with missing data skew the projection or rmsd is not a good metric in this case.

#### 3.1.2 RMSD of CA of Activation Loop, No Missing Residues

RMSD is carried out on the activation loop, which was previously identified as the highest average structural variance subsequence^3^. For the sake of comparison with the other methods upon the same segment, RMSD is calculated for the specific segment of all the conformations without any missing residues.

**Figure 3.5:**
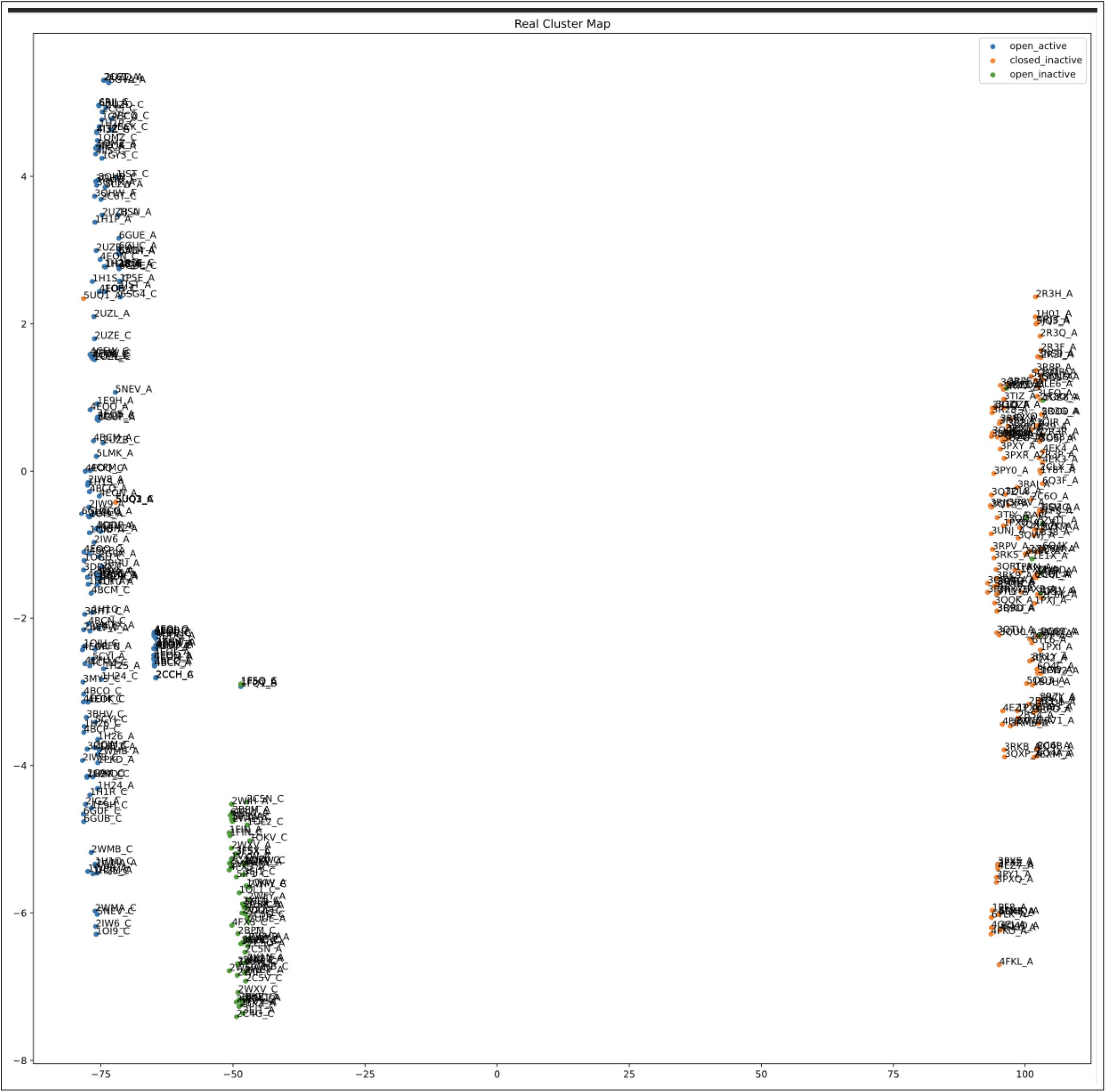
Real Cluster t-SNE projected map of RMSD of the activation loop

**Figure 3.6:**
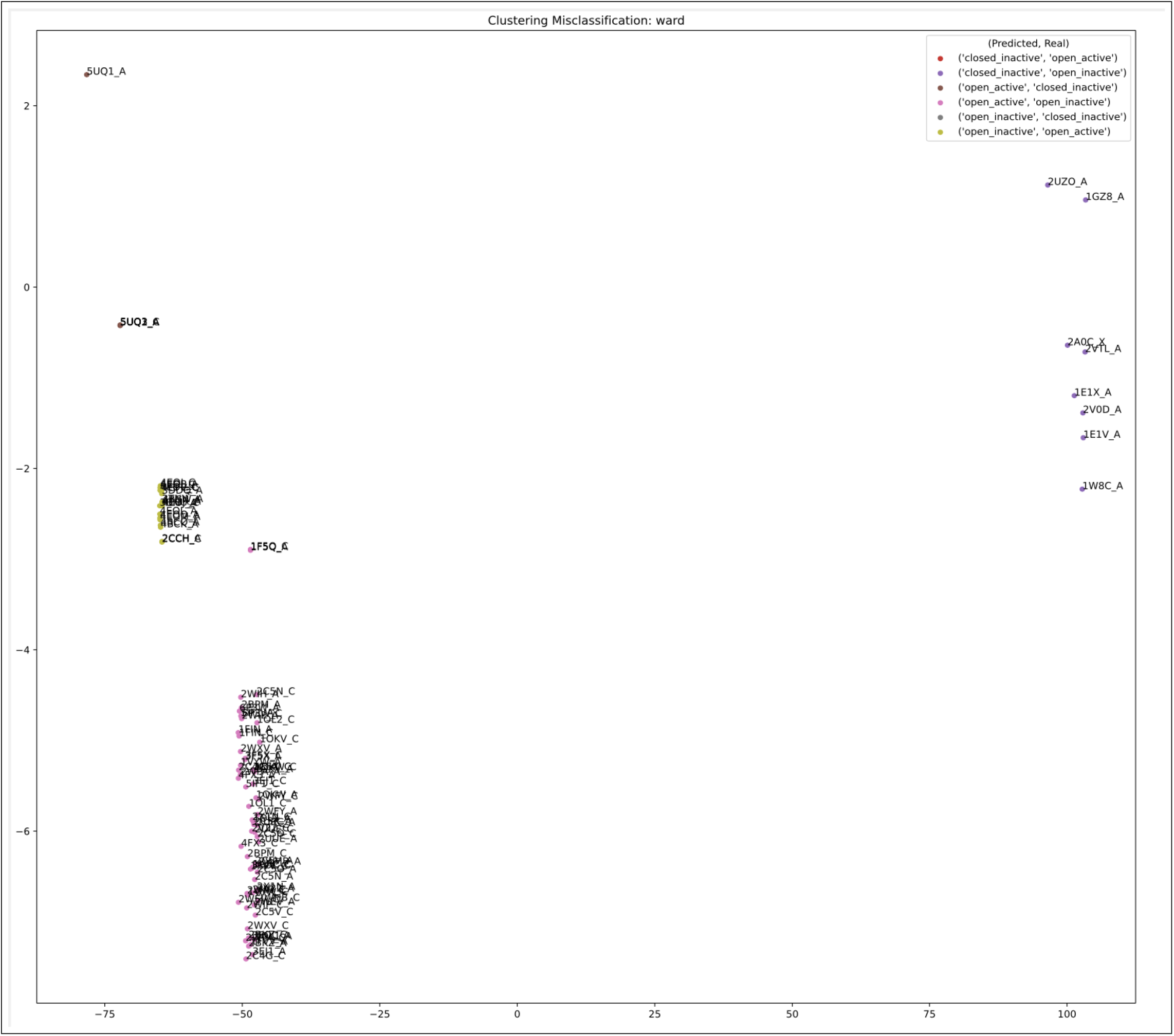
Misclassification Map of RMSD CA activation loop no missing residues

**Figure 3.7:**
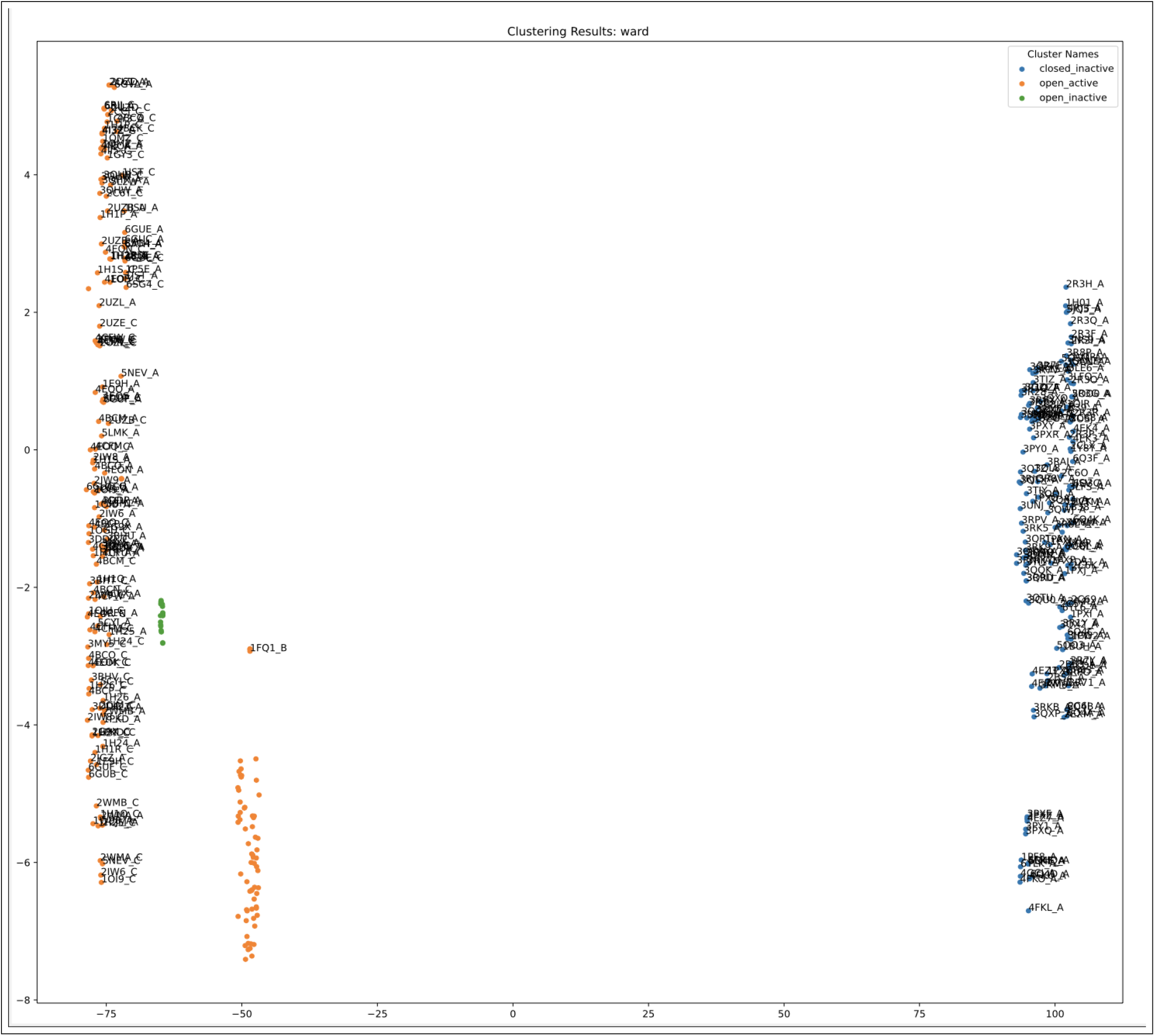
Clustering map of the ward’s algorithm. A surprising small enclave of open active is misclassified as open inactive

##### Statistics

**Table 3.3:**
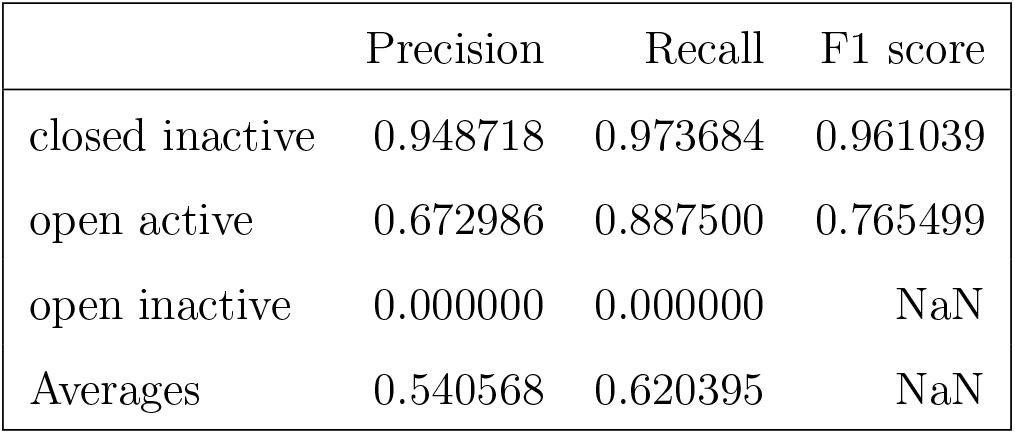
RMSD Activation Loop with no missing residues evaluation metrics

|  | Precision | Recall | F1 score |
| --- | --- | --- | --- |
| closed inactive | 0.948718 | 0.973684 | 0.961039 |
| open active | 0.672986 | 0.887500 | 0.765499 |
| open inactive | 0.000000 | 0.000000 | NaN |
| Averages | 0.540568 | 0.620395 | NaN |

**Table 3.4:** RMSD Activation Loop with no missing residues cross-classification table

|  | Real closed inactive | Real open active | Real open inactive |
| --- | --- | --- | --- |
| Predicted closed inactive | 148 | 0 | 8 |
| Predicted open active | 4 | 142 | 65 |
| Predicted open inactive | 0 | 18 | 0 |

##### Analysis

Although the clusters were well defined, due to large intra-cluster distances of open active, open inactive was completely misclassified. Potentially using a kernel projection, different distance metrics, different dendrogram cut, or a different clustering algorithm might work better.

Upon further inspection, 5UQ1 A and C, an outlier looks the same structurally as open active on pymol with only the abscence of TPO, despite being annotated as closed inactive. Furthermore, the small enclave of conformations mislabelled as open inactive are those which have very similar structures to 4EOJ A (open active) including 4EOJ A itself. This misclassification resulted in a whole group, open inactive misclassified and thus zero division error producing NaN for the F1 score. Despite the misclassification, the method produced heterogeneous clusters which shows that high variance regions were correctly identified and correctly identify each conformation too.

### 3.2 t-SNE Based Map Clustering results

The plots below describe one trials of the t-SNE Map Clustering^4^ of the specific experiment for visualization, based on the multi-trial statistics these plots should remain relatively consistent. The statistics, on the other hand, are of the 20 trials of the specific experiment since t-SNE is contains of random initialization.

#### 3.2.1 Complete t-SNE Based Map based clustering CA

Now we test the ability of the t-SNE map based clustering against that of the full RMSD without the high variance method and using imputations for the missing values. Also since both of the plots are one dimensional, y-axis is imputed with 1s and labels are removed due to overcrowding so that the clusters can be visualized.

**Figure 3.8:**
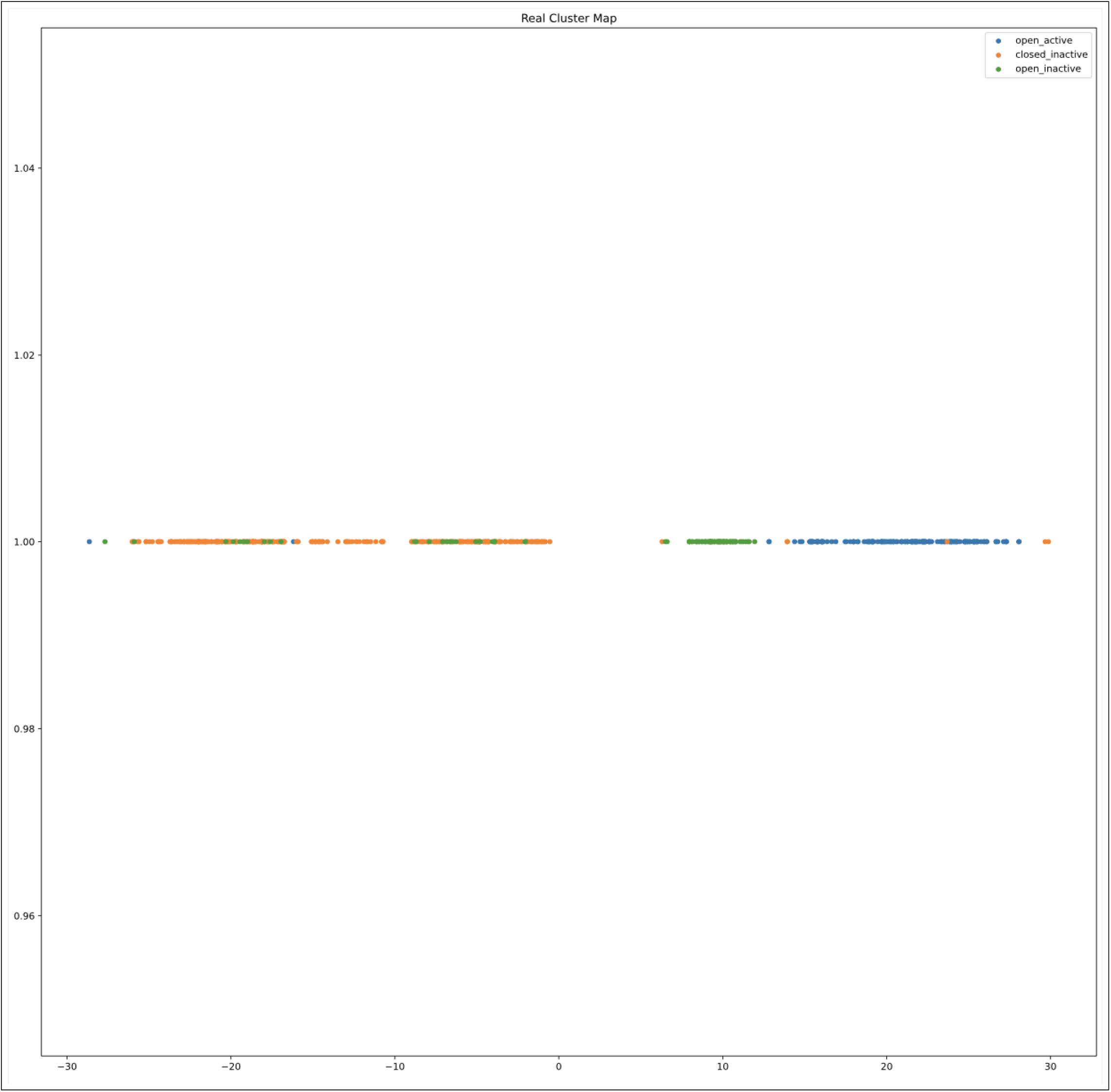
Real Cluster t-SNE projected map of t-SNE based clustering of the Activation Loop, No missing residues

**Figure 3.9:**
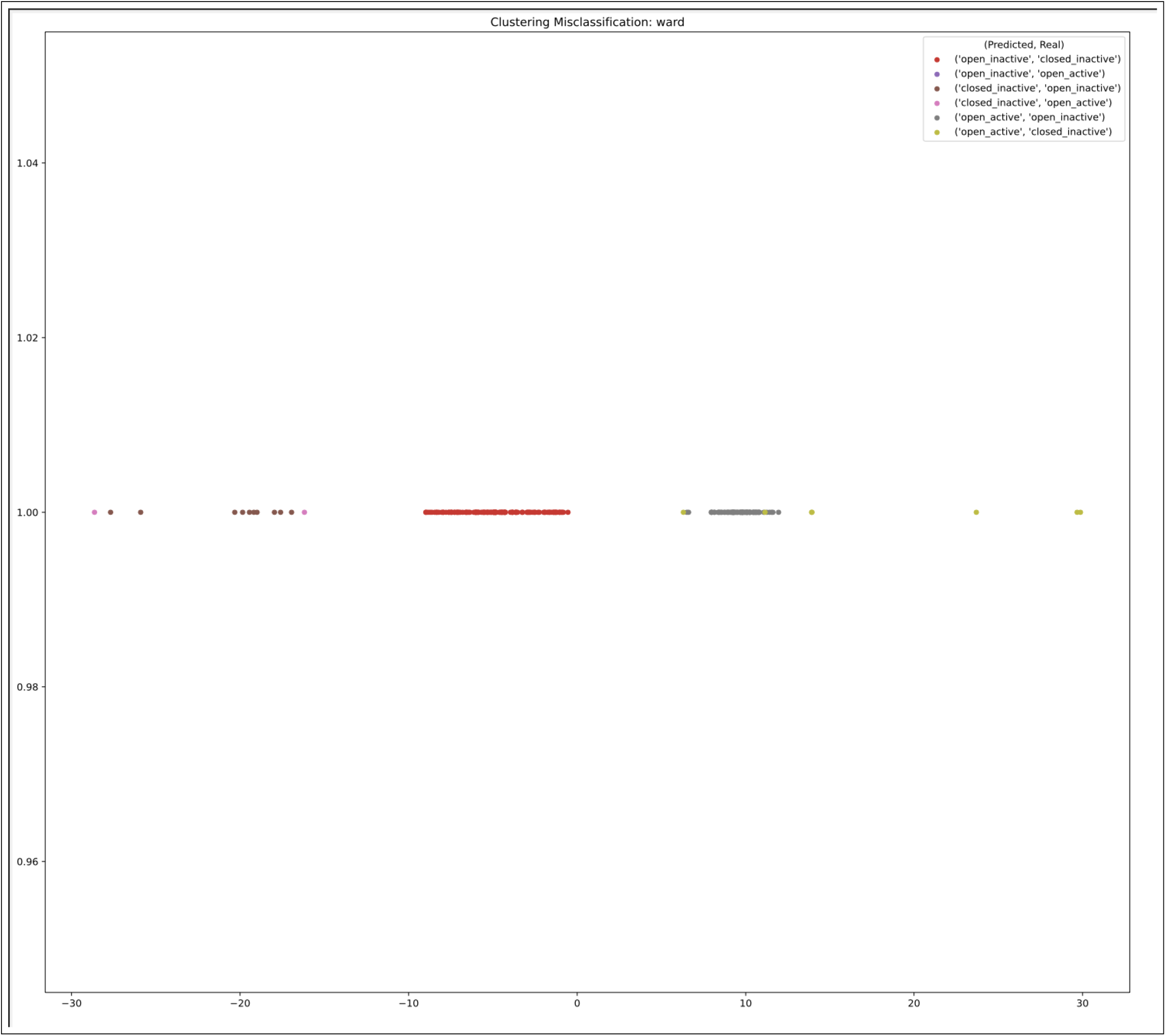
Misclassification of t-SNE Based map of full sequence

**Figure 3.10:**
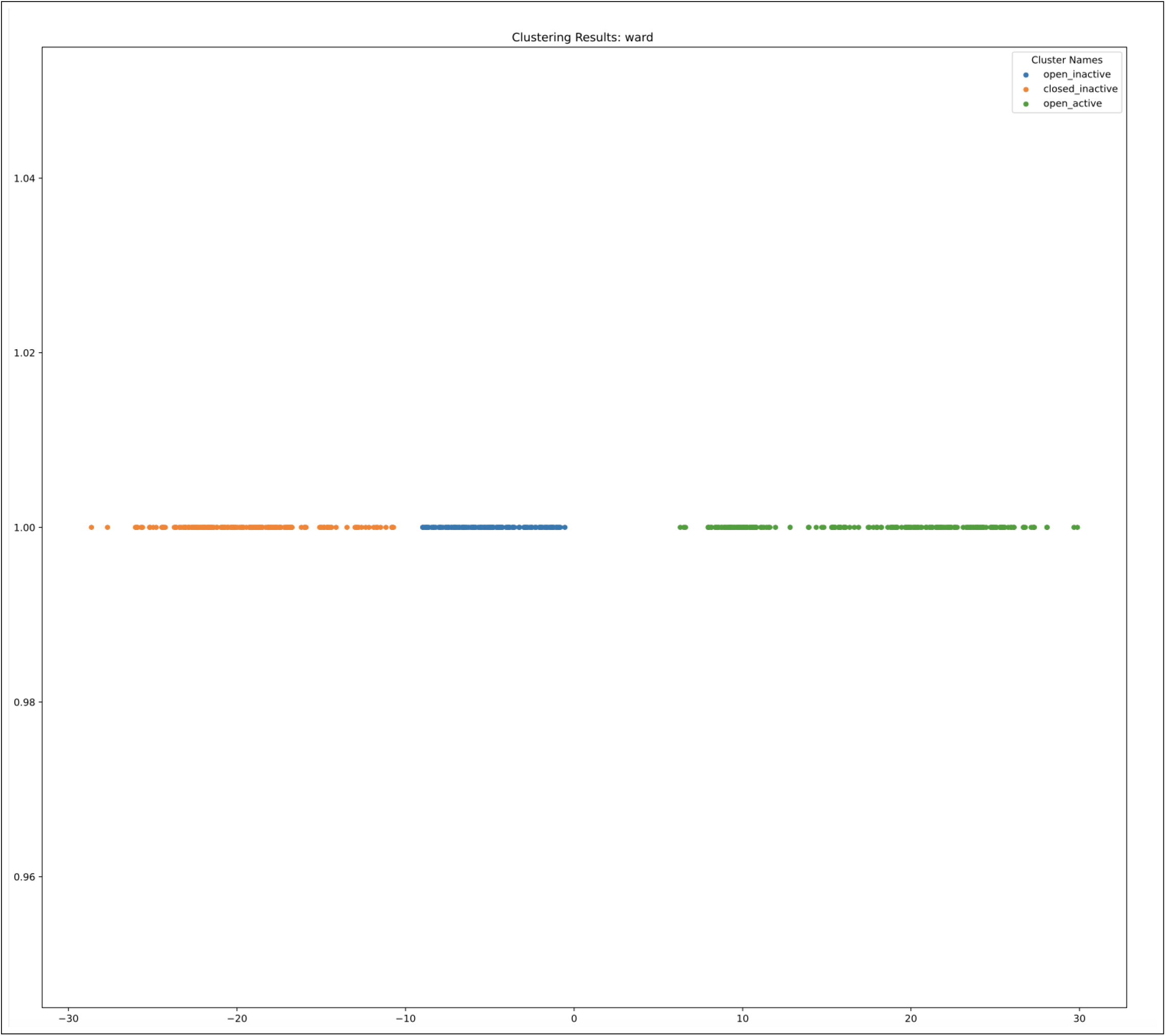
Clustering Map of t-SNE Based map clustering of full sequence using ward’s algorithm

**Figure 3.11:**
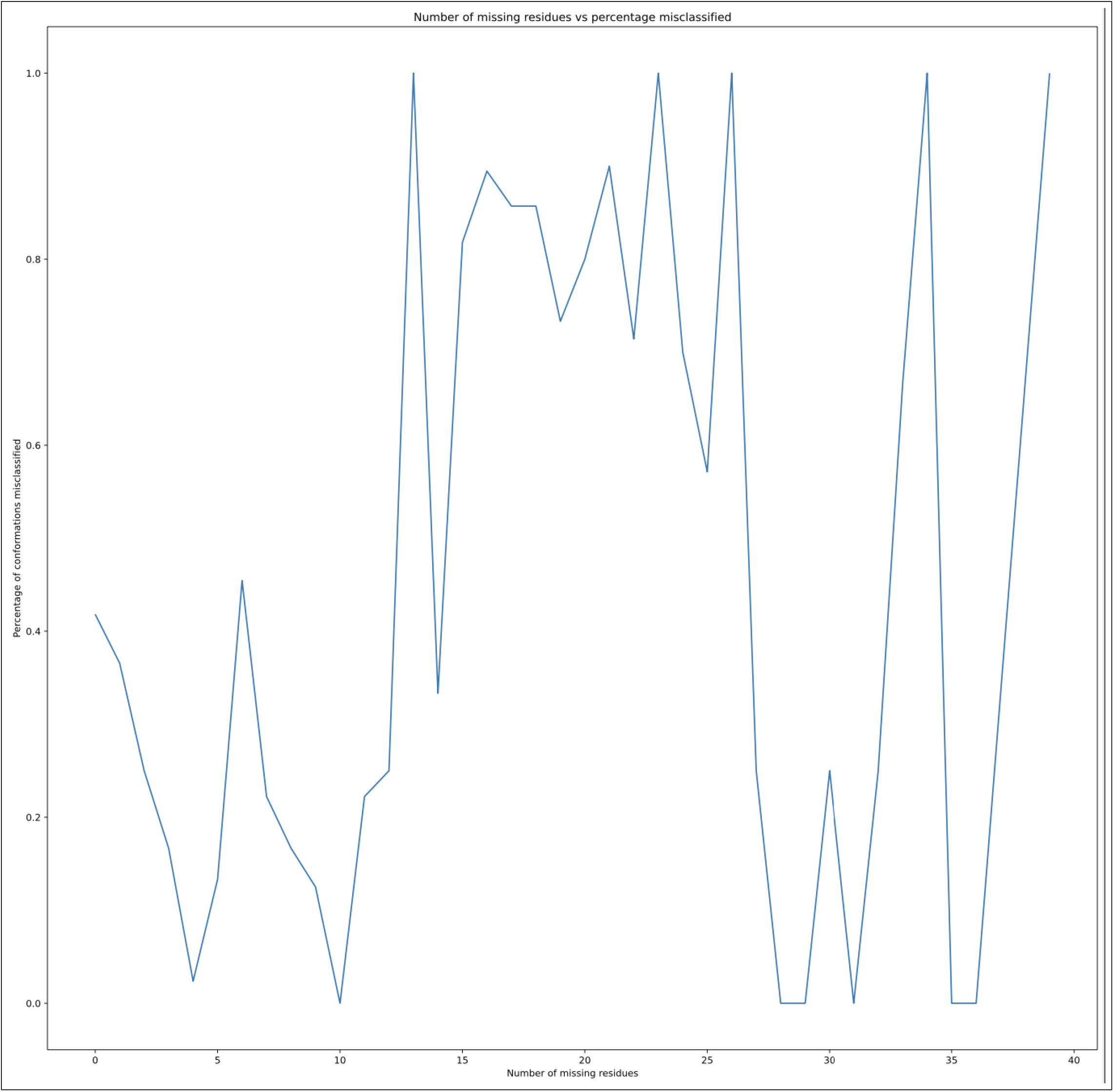
Misclassification vs Missing Residues

##### Statistics

**Table 3.5:** complete t-SNE based map trials

|  | Precision | Recall | F1 score |
| --- | --- | --- | --- |
| Trial 1 | 0.584171 | 0.582976 | 0.559980 |
| Trial 2 | 0.575304 | 0.574047 | 0.550408 |
| Trial 3 | 0.573939 | 0.558055 | 0.536026 |
| Trial 4 | 0.580932 | 0.579366 | 0.555101 |
| Trial 5 | 0.576374 | 0.575600 | 0.553737 |
| Trial 6 | 0.567864 | 0.549631 | 0.526155 |
| Trial 7 | 0.573049 | 0.571990 | 0.548872 |
| Trial 8 | 0.565355 | 0.547560 | 0.525323 |
| Trial 9 | 0.577491 | 0.576454 | 0.553656 |
| Trial 10 | 0.572089 | 0.556851 | 0.533822 |
| Trial 11 | 0.575575 | 0.575251 | 0.552178 |
| Trial 12 | 0.575733 | 0.575600 | 0.553278 |
| Trial 13 | 0.583085 | 0.581773 | 0.558354 |
| Trial 14 | 0.581456 | 0.581773 | 0.558655 |
| Trial 15 | 0.579057 | 0.578861 | 0.556437 |
| Trial 16 | 0.572809 | 0.556851 | 0.534318 |
| Trial 17 | 0.578439 | 0.568536 | 0.545436 |
| Trial 18 | 0.583035 | 0.582122 | 0.560065 |
| Trial 19 | 0.580831 | 0.579715 | 0.556818 |
| Trial 20 | 0.583529 | 0.582976 | 0.559514 |

**Table 3.6:** complete t-SNE based map trials, Average Metrics

|  | Precision | Recall | F1 score |
| --- | --- | --- | --- |
| open inactive | 0.117027 | 0.162500 | 0.135928 |
| closed inactive | 0.931869 | 0.566787 | 0.704379 |
| open active | 0.682121 | 0.986111 | 0.806414 |
| Averages | 0.577006 | 0.571799 | 0.548907 |

**Table 3.7:** complete t-SNE based map trials, Average Cross Classification Statistics

|  | Real open inactive | Real closed inactive | Real open active |
| --- | --- | --- | --- |
| Predicted open inactive | 14.95 | 113.4 | 0.00 |
| Predicted closed inactive | 9.20 | 157.0 | 2.25 |
| Predicted open active | 67.85 | 6.6 | 159.75 |

##### Analysis

The misclassification percentage vs the missing residues graph resembled that of the RMSD for the similar full chain without discarding samples with missing residues. This technique; however, as opposed to the RMSD with similar parameters, the t-SNE based map did slightly better and created more homogeneous and more distinct clusters. A similar occurrence with open active and open inactive group, it mostly suffered from the shift of the clustering towards the left. Perhaps a change of clustering metric, dendrogram cut, or method would give a higher F1 score.

#### 3.2.2 t-SNE Based Map Clustering of CA of Activation Loop, No Missing Residues

This experiment was run on the highest variance subsequence derived from the technique as mentioned above, after discarding all of the conformations with more than 1 residue missing within the subsequence in focus.

**Figure 3.12:**
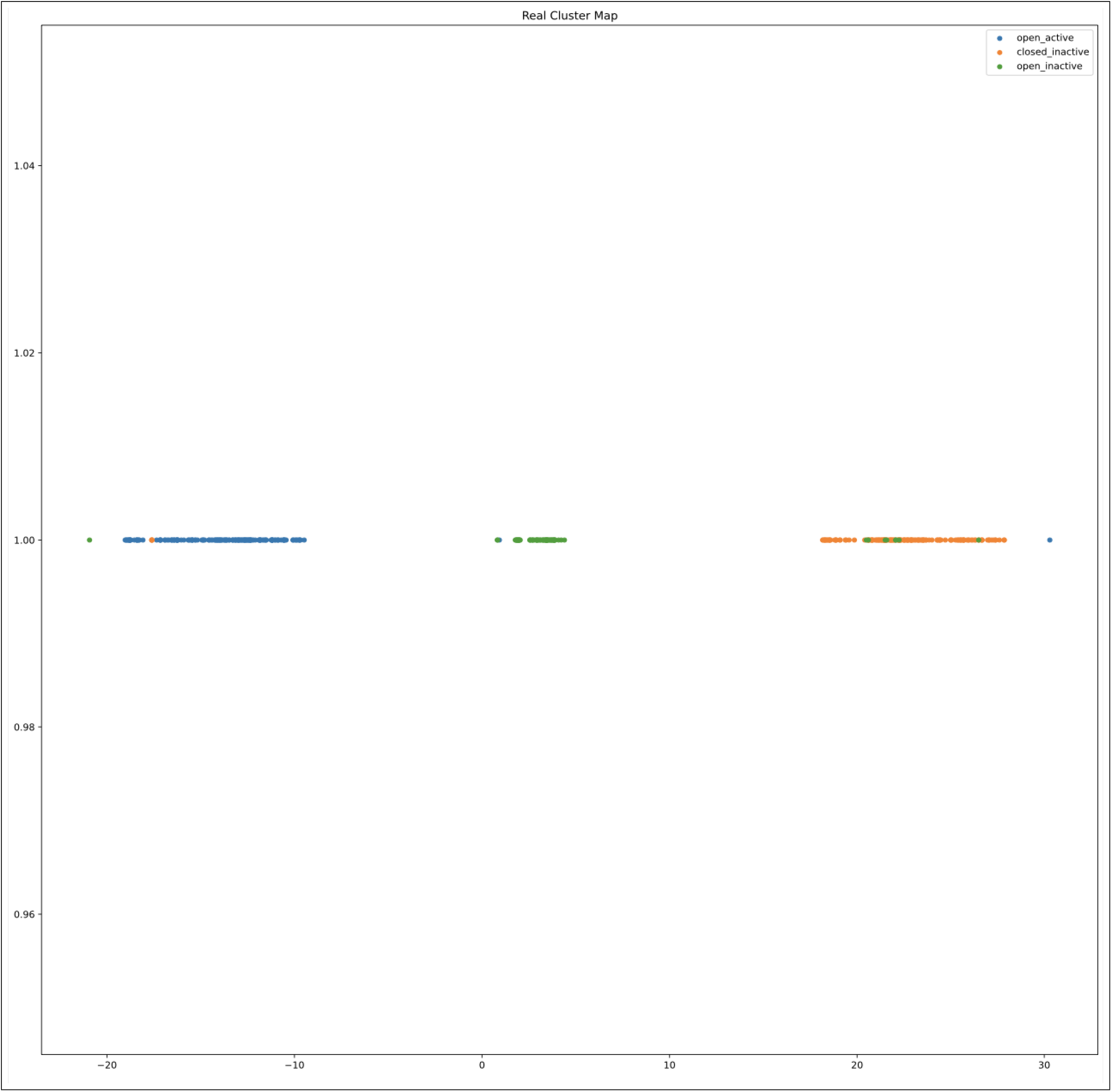
Real Cluster t-SNE projected map of t-SNE based clustering of the whole sequence

**Figure 3.13:**
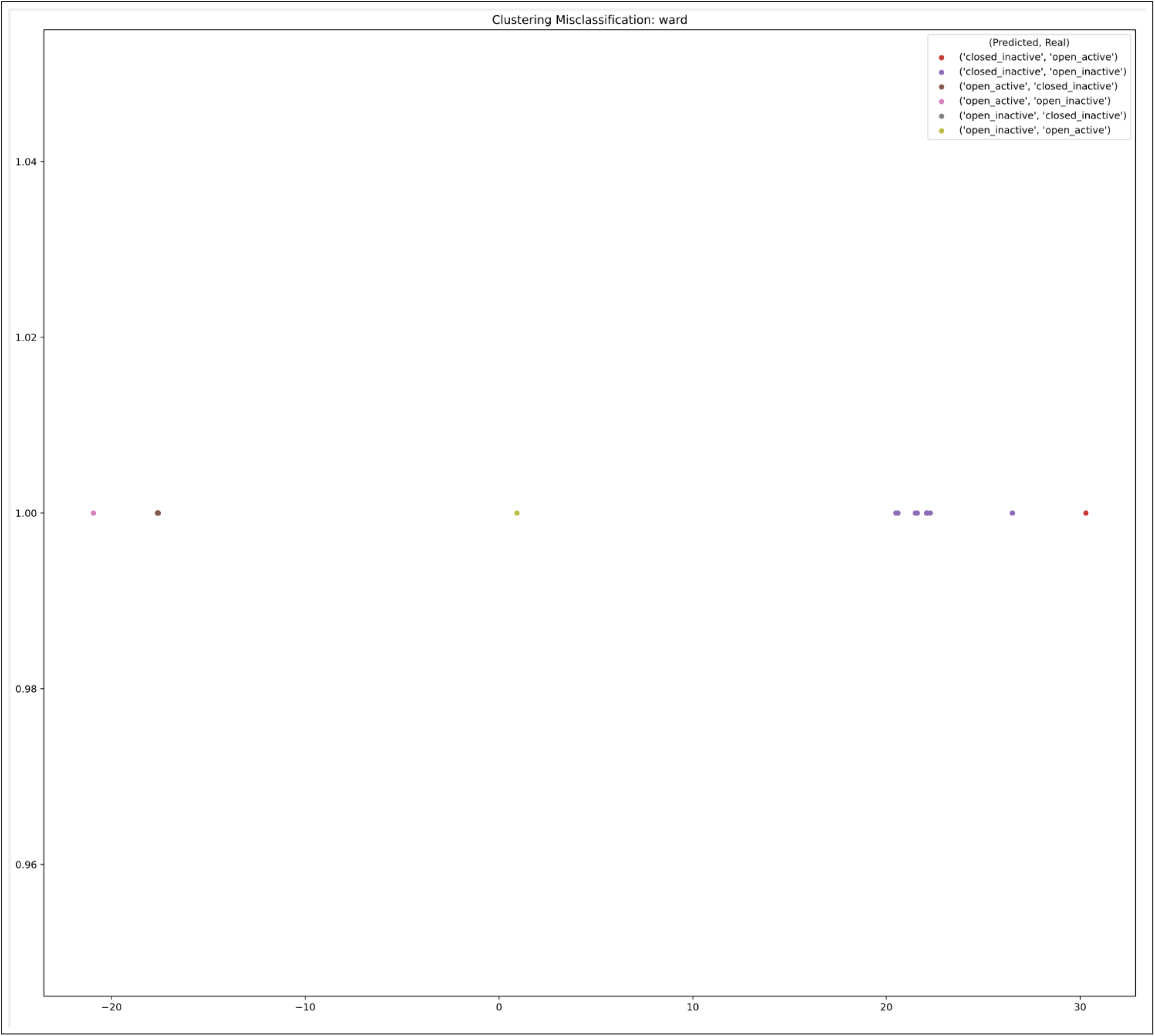
Misclassification of t-SNE based map clustering of Activation loop, No missing residues

**Figure 3.14:**
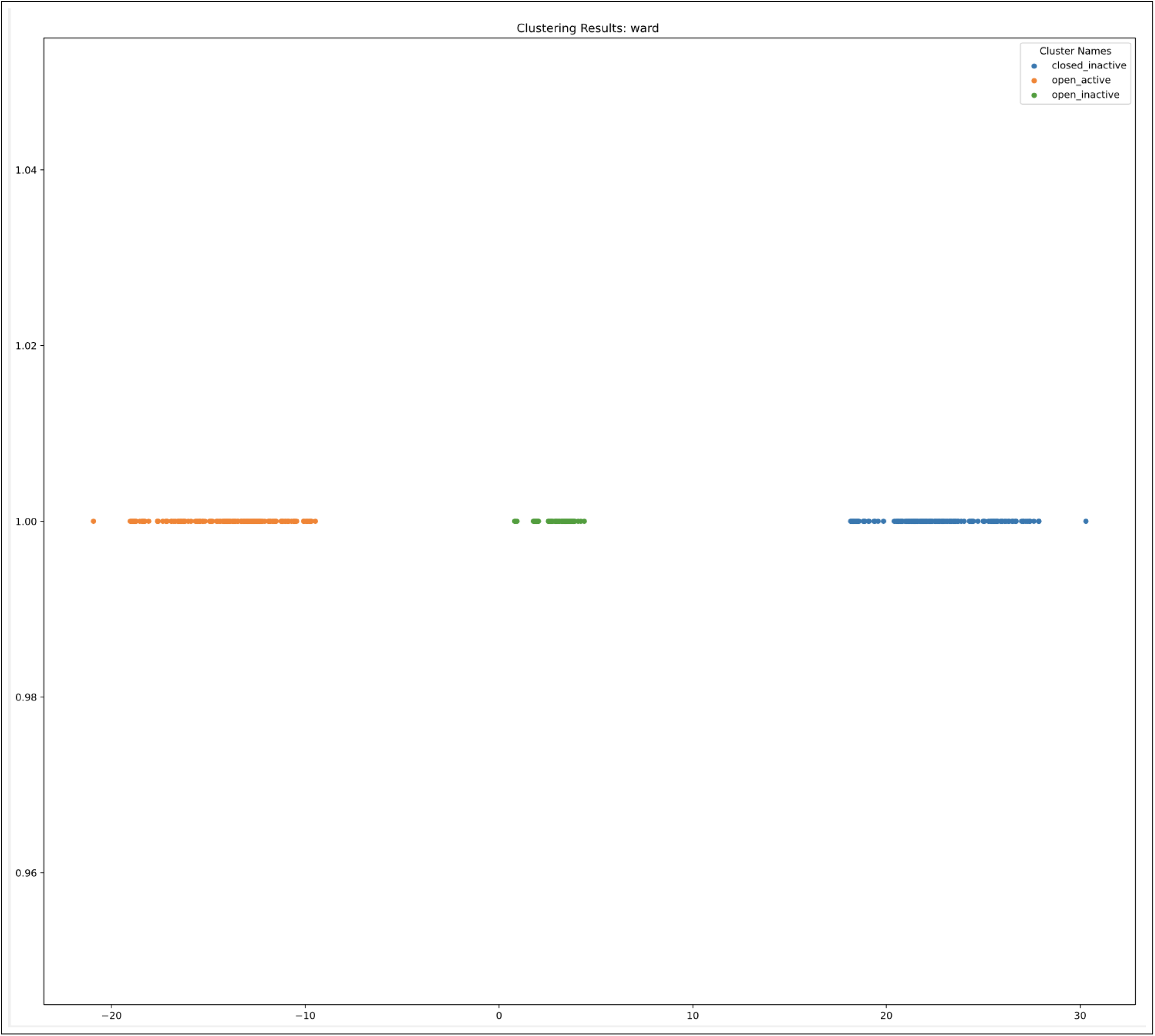
Clustering Map of t-SNE Based map clustering of activation loop using ward’s algorithm, No missing residues

##### Statistics

**Table 3.8:** Activation Loop with no missing residues, t-SNE Based Map Clustering trials

|  | Precision | Recall | F1 score |
| --- | --- | --- | --- |
| Trial 1 | 0.962589 | 0.948339 | 0.954828 |
| Trial 2 | 0.957627 | 0.925218 | 0.938274 |
| Trial 3 | 0.945022 | 0.928461 | 0.935884 |
| Trial 4 | 0.954280 | 0.939676 | 0.946329 |
| Trial 5 | 0.949099 | 0.930364 | 0.938615 |
| Trial 6 | 0.932497 | 0.921592 | 0.926599 |
| Trial 7 | 0.946198 | 0.937523 | 0.941568 |
| Trial 8 | 0.946597 | 0.935220 | 0.940486 |
| Trial 9 | 0.953146 | 0.925798 | 0.937264 |
| Trial 10 | 0.941466 | 0.909906 | 0.922616 |
| Trial 11 | 0.941744 | 0.930325 | 0.935497 |
| Trial 12 | 0.939410 | 0.926197 | 0.932202 |
| Trial 13 | 0.955556 | 0.941689 | 0.947995 |
| Trial 14 | 0.963440 | 0.941290 | 0.950905 |
| Trial 15 | 0.955269 | 0.930364 | 0.941002 |
| Trial 16 | 0.951299 | 0.937303 | 0.943682 |
| Trial 17 | 0.959327 | 0.939387 | 0.948191 |
| Trial 18 | 0.942283 | 0.928680 | 0.934812 |
| Trial 19 | 0.943051 | 0.923894 | 0.932314 |
| Trial 20 | 0.900812 | 0.897148 | 0.898896 |
| Average | 0.944180 | 0.927715 | 0.934945 |

**Table 3.9:** Activation Loop with no missing residues, Average Metrics

|  | Precision | Recall | F1 score |
| --- | --- | --- | --- |
| closed_inactive | 0.931295 | 0.953289 | 0.942127 |
| open_active | 0.954195 | 0.974375 | 0.964127 |
| open_inactive | 0.947051 | 0.855479 | 0.898581 |
| Averages | 0.944180 | 0.927715 | 0.934945 |

**Table 3.10:** Activation Loop with no missing residues, Average Cross Classification Statistics

|  | Real closed inactive | Real open active | Real open inactive |
| --- | --- | --- | --- |
| Predicted closed inactive | 145.50 | 1.80 | 8.65 |
| Predicted open active | 5.55 | 156.35 | 1.80 |
| Predicted open inactive | 0.95 | 1.85 | 62.55 |

##### Analysis

The results vary by initialization, however, probably by only a few cross classification of samples at at time. Our statistical metrics: Precision, Recall, and F1 score are consistently high for each run. The clusters are more distinct as compared to the RMSD method and are to an extent homogeneous in the real cluster map. The problem between closed inactive predicted as open active at 5.55 average cross classification is solved. In general the results are consistent across the trials. Beyond this the clusters are seen to be consistent with the annotated groupings after the t-SNE map. The method proved to do well all around given full data on the activation loop.

### 3.3 Dihedral Based Clustering

In this experiment we use the PSS software to do dihedral based clustering^5^ of the structures of all the samples without missing residues (Gaillard et al. 2013).

**Figure 3.15:**
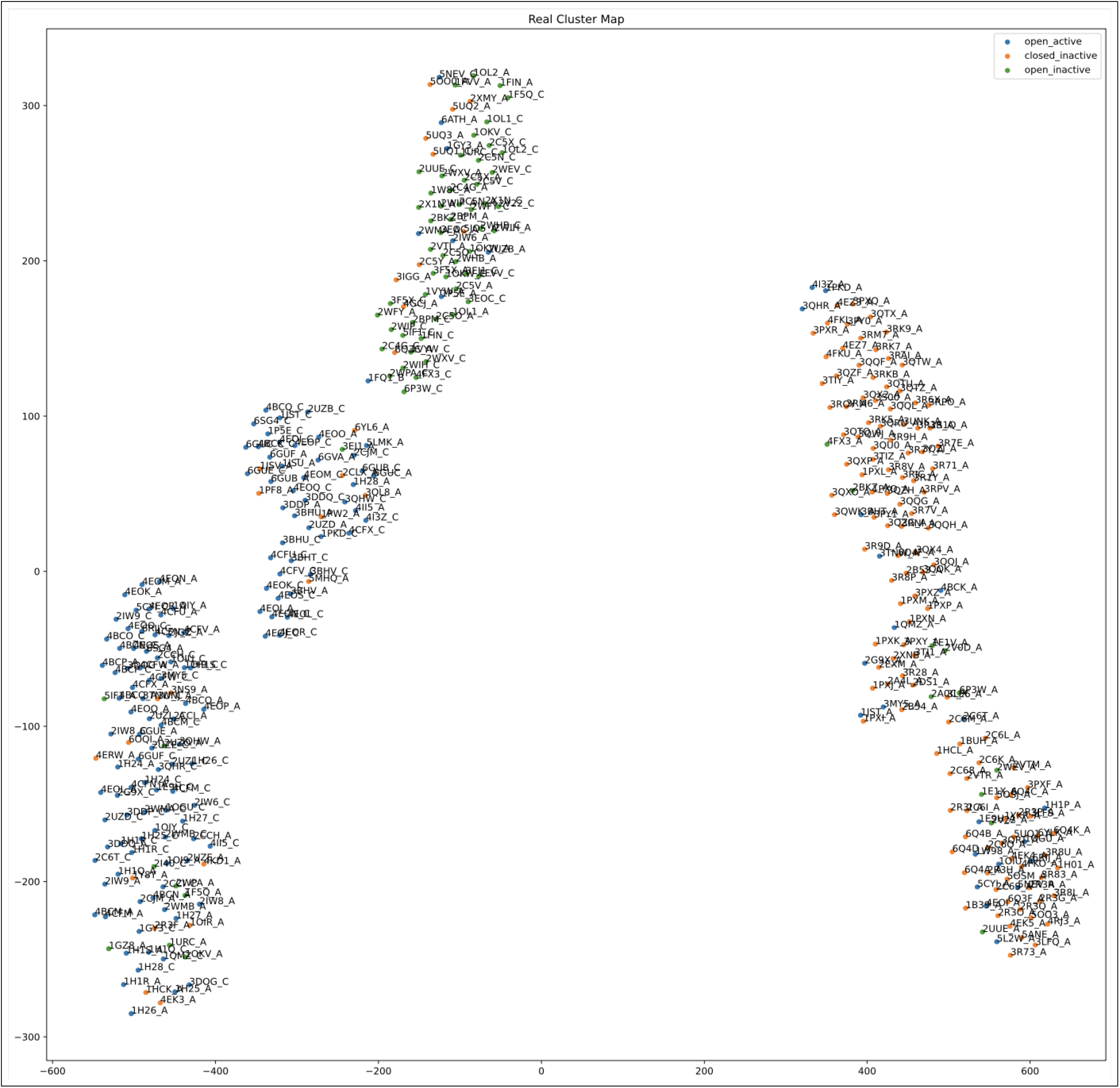
Real Cluster of Dihedral-Based clustering of activation loop

**Figure 3.16:**
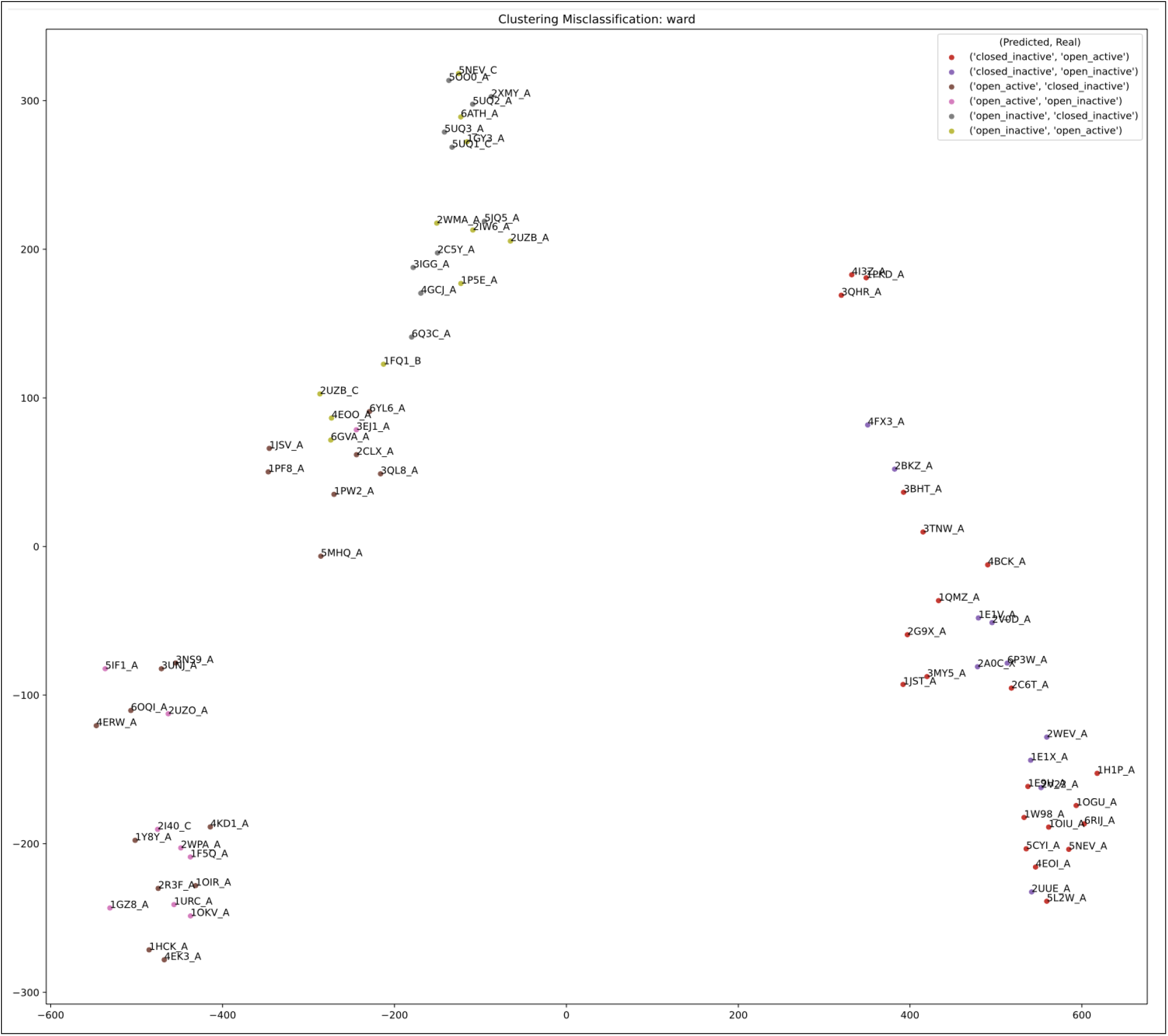
Misclassification of Dihedral-Based clustering of activation loop

**Figure 3.17:**
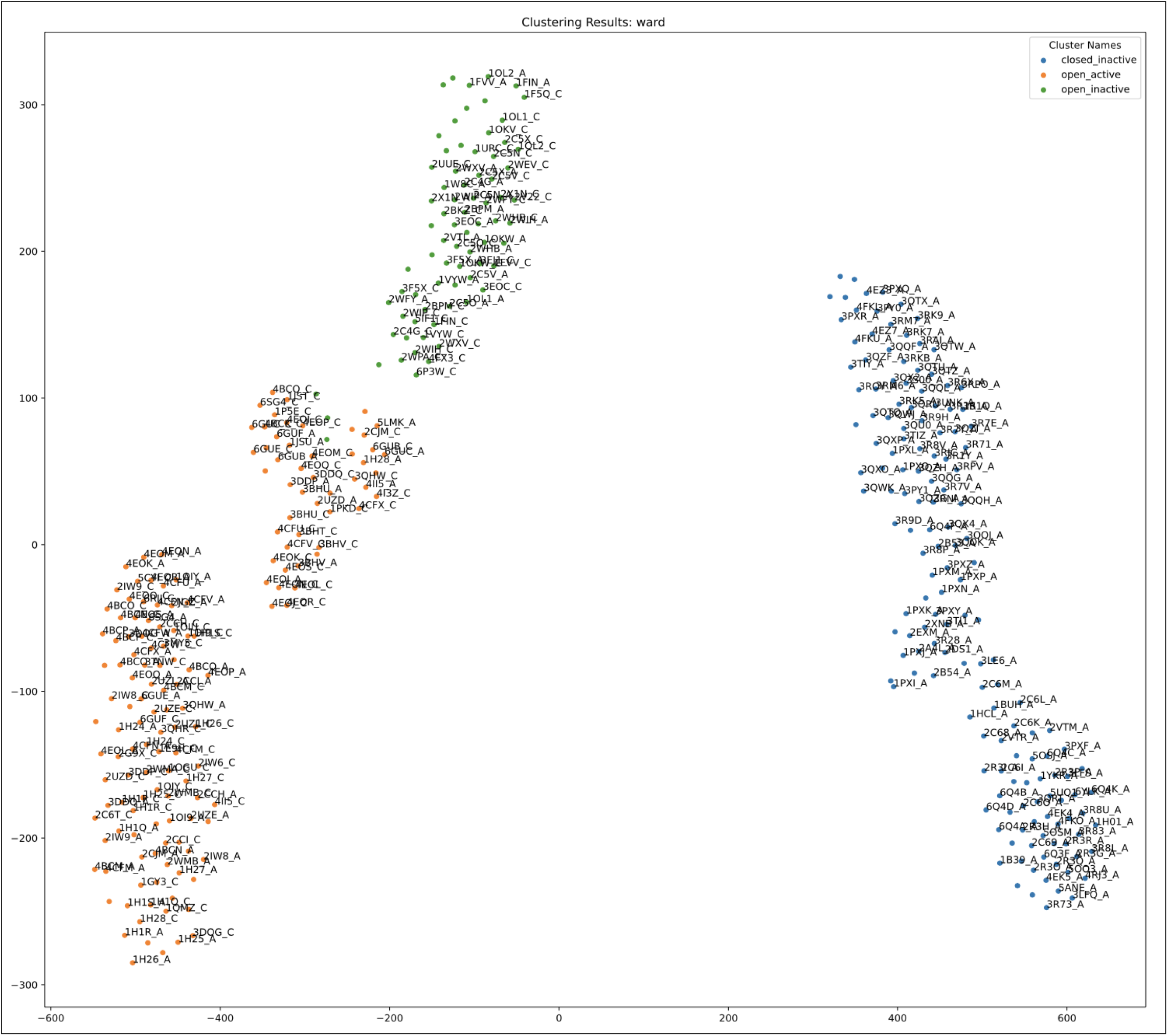
Clustering Map of Dihedral-Based clustering of activation loop

#### Statistics

**Table 3.11:** Dihedral-Based Clustering: Activation Loop with no missing residues, Statistics Metrics

|  | Precision | Recall | F1 score |
| --- | --- | --- | --- |
| closed inactive | 0.800000 | 0.821192 | 0.810458 |
| open active | 0.831169 | 0.800000 | 0.815287 |
| open inactive | 0.720000 | 0.739726 | 0.729730 |
| Averages | 0.783723 | 0.786973 | 0.785158 |

**Table 3.12:** Dihedral-Based Clustering: Activation Loop with no missing residues, Cross Classification Statistics

|  | Real closed inactive | Real open active | Real open inactive |
| --- | --- | --- | --- |
| Predicted closed inactive | 124 | 21 | 10 |
| Predicted open active | 17 | 128 | 9 |
| Predicted open inactive | 10 | 11 | 54 |

#### Analysis

Unlike the RMSD of the activation loop, which sacrificed precision for recall or the F1 score of one group for another, the dihedral based matrix is relatively more heterogeneous throughout its clusters as compared to the previous two methods. The dihedral-based classification method does well all-around since it gives insight into shape and structure within a conformation. However, clusters were not as distinct as the t-SNE based map and the lack of homogeneity causes consistent misclassification throughout the groups. Despite the heterogeneity the method does perform decently due to the slightly higher concentrations of each group in each cluster.

## Conclusion

### 4.1 Summary of methods

The RMSD matrix and T-SNE based map clustering both gave consistent results and clustered the conformations well. The RMSD based clustering although less costly and more stable, occasionally was thwarted by the high intra-clusteral distances although the conformations were well clustered.

The T-SNE based map, on the other hand, despite, being a bit more costly requiring multiple trials and input parameters like perplexity and high variance region qualification percentiles, produced much more heterogeneous distinct clusters based on position instead of relative position as RMSD. This showed fruition in out performing both RMSD and Dihedral based clustering in the statistical metrics. Some part of the distinctness of the clusters can be attributed to extra t-SNE projection with the high perplexity which with the Kullback-Leiber divergence penalization, created clear neighborhoods for each cluster. In both RMSD and t-SNE clustering, the methods applied on specific regions with high structural variance both performed much better as it amplified the differences previously suppressed by the other parts of the sequence which were pair-wise similar.

The t-SNE based map clustering, in general provides user flexibility— able to control for certain high variance regions, to pick and choose each segment and include them as features and has consistently out performed the other methods. It has some variable parameters like clustering perplexity, initialization, and high variance percentiles, which can, however, be generalized since high perplexity will always form distinct clusters, multiple runs can give an average result, and high variance regions will be distinct on the plot for the user to create cuts.

In the end, RMSD is straight-forward and robust, however, occasionally doesn’t create homogeneous and distinct clusters, given that we don’t know the actual groupings in actual applications, can be problematic. Aside from the few weaknesses, empowered with identified high variance regions, it can perform well.

Dihedral-based is straight-forward and robust as well, giving a well balanced result. On the other hand it creates more heterogeneous clusters of the conformations compared to the two previous methods, often mixing different groups in the same cluster, this is penalized in the statistical metrics.

### 4.2 Discussion

#### 4.2.1 Finding High Variance Regions

Currently, the method of detecting high variance regions requires similar numbering and residues between conformations. However, for a data set of conformations with more heterogeneous structures, this method can be applied after doing some sequence and structural alignment.

On top of this, despite being an efficient map in the case of clustering CDK2 proteins, in order to deal with symmetrical substructures which have the same average coordinates a more sophisticated map should be devised. A proposition of using the current methods to address the problem is to again cut the subsequence into half and use each half as a feature instead as all together, however, this requires inspection of the physical structure which defeats the goal of automation.

#### 4.2.2 Clustering and Mapping Methods

The t-SNE based map clustering is very rough and many more optimizations can be built upon it. For example currently the imputation method is very primitive only taking the averages of the coordinates of the same indices in other conformations this can lead to the formation of an artificial cluster made from the averages of all the groups. Perhaps a method can be formulated which might take into account the coordinates of the neighboring residues in a weighted manner to impute a more precise result.

As seen throughout both of the t-SNE based map clustering and RMSD matrix, despite the correct clustering of the methods, intra-cluster distances tend to hamper with the results causing a proportion of the correctly clustered group to be misclassified. A different distance metrics or clustering techniques could be used to address this issue.

## Footnotes

1 Results obtained from https://github.com/yao-creative/cdk2_human_classification.git

2 refer to section 2.4

3 refer to section 2.3

4 refer to section 2.6

5 refer to section 2.5

## Notes

https://github.com/yao-creative/cdk2_human_classification

### Competing Interest Statement

The authors have declared no competing interest.

